# The Hidden Costs of Climate Change Tracking: Climate Velocity, Movement Energetics, and Connectivity in European Protected Areas

**DOI:** 10.64898/2026.04.20.719608

**Authors:** Gavin Stark, Jeremy Dertien, Nikolaj Rauff Poulsen, Emilio Berti, Andres Camilo Mármol Guijarro, Magali Weissgerber, Néstor Fernández, Henrique M. Pereira

**Author notes:** Corresponding author: Gavin Stark; mailing address: German Center of Integrative Biodiversity Research (iDiv), Puschstraße 4, 04103 Leipzig, Germany.

## Abstract

Rapid climate change is reducing the capacity of protected areas (PAs) to conserve biodiversity, but exposure metrics alone do not show whether species can reach areas where suitable climates persist. We developed a climate-informed connectivity framework integrating climate velocity, PA climatic residence time, PA size, and functional connectivity based on energetics-informed resistance surfaces from species distribution models.

Using high-resolution climate projections and omnidirectional connectivity modelling across Europe, we show that climate-tracking opportunities are more limited and spatially uneven than structural connectivity alone suggests. Small, climate-exposed PAs are especially vulnerable because they provide little internal climatic buffering and are often embedded in landscapes with low movement feasibility, whereas larger and more climatically stable PAs are more often situated in landscapes that can support redistribution. These findings provide a spatially explicit basis for restoration and conservation planning to maintain the functionality of PA networks under future climate change.

**Teaser:** Many European protected areas are too isolated, climate-exposed, and energy-constrained to support climate-tracking connectivity.

## Introduction

Protected areas (PAs) remain central to biodiversity conservation, yet rapid climate change is increasingly undermining their ability to secure long-term persistence (Cui et al., 2025). Conservation planning has historically been grounded in contemporary species distributions and present-day ecological conditions, often without explicitly accounting for how climate change will reorganize habitats and communities through time (Evans, 2012; *Natura 2000*, 2009; Rodríguez-Rodríguez & Martínez-Vega, 2022). This mismatch between largely static protection and dynamic climate exposure is already evident across Europe, where shifts in temperature and precipitation are reshaping species ranges, phenology, and ecosystem functioning, threatening the climatic integrity of many sites designated to safeguard biodiversity (Araújo et al., 2011; Dobrowski et al., 2021; Hoffmann et al., 2019; Hülber et al., 2020; Johnston et al., 2013; Kelly et al., 2025; Sonntag & Fourcade, 2022; Wessely et al., 2017). As a result, the long-term value of PAs increasingly depends on whether species can move beyond their boundaries as local conditions become unsuitable (Dobrowski et al., 2021).

Assessing whether biodiversity can keep pace with climate change requires metrics that link the rate of environmental change to the capacity of organisms to track suitable conditions (Sandel et al., 2011; Thomas et al., 2004). Estimates of climate velocity provide such a link by translating climatic change into the geographic distances species must traverse to remain within similar climatic environments (García Molinos et al., 2019; Loarie et al., 2009; Sandel et al., 2011). Where climate velocity is high, climatic residence times are short, constraining the window for local persistence before relocation becomes necessary (Brito-Morales et al., 2018; Pacifici et al., 2015; Thomas et al., 2004; Urban et al., 2012). Across Europe, this exposure is uneven: intensively modified lowlands are projected to undergo rapid climatic turnover, whereas topographically complex mountain systems experience slower change and greater microclimatic heterogeneity, increasing their likelihood of functioning as climatic refugia (Cimatti et al., 2025; Cui et al., 2025; Dragonetti et al., 2024; Heikkinen et al., 2020; Loarie et al., 2009). This asymmetry creates a directional imperative for movement from fast-changing lowlands toward more climatically stable regions (Cimatti et al., 2025).

In this context, ecological connectivity is critical for enabling climate-driven range shifts, sustaining gene flow, and facilitating movement as conditions deteriorate (da Silva et al., 2024; Hilty et al., 2020; Littlefield et al., 2017, 2019; McGuire et al., 2016). This need is increasingly reflected in policy, from the EU Biodiversity Strategy for 2030 and proposals for a Trans-European Nature Network that emphasize ecological corridors, to the Kunming–Montreal Global Biodiversity Framework, whose Target 2 links large-scale ecosystem restoration explicitly to the recovery of ecological integrity and connectivity (Hermoso et al., 2022; Lapin et al., 2025; Moreira et al., 2024; Stephens, 2023). However, many connectivity assessments remain largely static and structural, emphasizing habitat continuity or present-day overlap without explicitly accounting for climate velocity or the need to connect climate-exposed PAs to long-term refugia (Keeley et al., 2018; Lapin et al., 2025; Nuñez et al., 2013). Consequently, corridors may appear connected in principle yet fail to support climate-adaptive movement in practice (Heller & Zavaleta, 2009; Reside et al., 2018).

A further limitation is the frequent assumption that movement is feasible wherever habitat continuity exists (Correa Ayram et al., 2016). Such approaches often overlook energetic and physiological constraints imposed by terrain, land use, vegetation structure, and thermal environments—factors that can slow dispersal, decrease reproductive success (fitness), increase mortality risk, and ultimately prevent populations from tracking shifting climates over long distances (Benoit et al., 2020; Berti et al., 2025; Briscoe et al., 2023; Cordes et al., 2025; Klarevas-Irby et al., 2025; Pontzer, 2016; Shepard et al., 2013; Stark et al., 2023). Ignoring these constraints risks identifying corridors that are structurally connected yet functionally inaccessible (Berti et al., 2022; Doherty & Driscoll, 2018; Shepard et al., 2013). Despite growing recognition of these issues, continental-scale connectivity assessments have rarely integrated climate exposure, climatic persistence, and the energetic feasibility of movement within a single framework.

Here, we develop a climate-informed connectivity framework that links protected-area exposure to climate change with the physiological feasibility of animal movement (Fig. 1). We combine climate velocity to identify PAs at risk of rapid climatic turnover, metrics of climatic stability to delineate potential long-term refugia, and movement energetics to evaluate whether dispersal between them is realistic rather than merely structurally possible. Applying this framework across Europe, we assess whether climate-exposed PAs are functionally connected to climatically stable regions, including those associated with topographically complex landscapes. By incorporating energetic constraints into climate-adaptive connectivity planning, our approach moves beyond habitat-only models and provides an operational basis to inform the planning of the Trans-European Nature Network, national restoration strategies, target 2 of the Kunming-Montreal Global Biodiversity Framework, and long-term conservation effectiveness under accelerating climate change.

**Figure 1.**
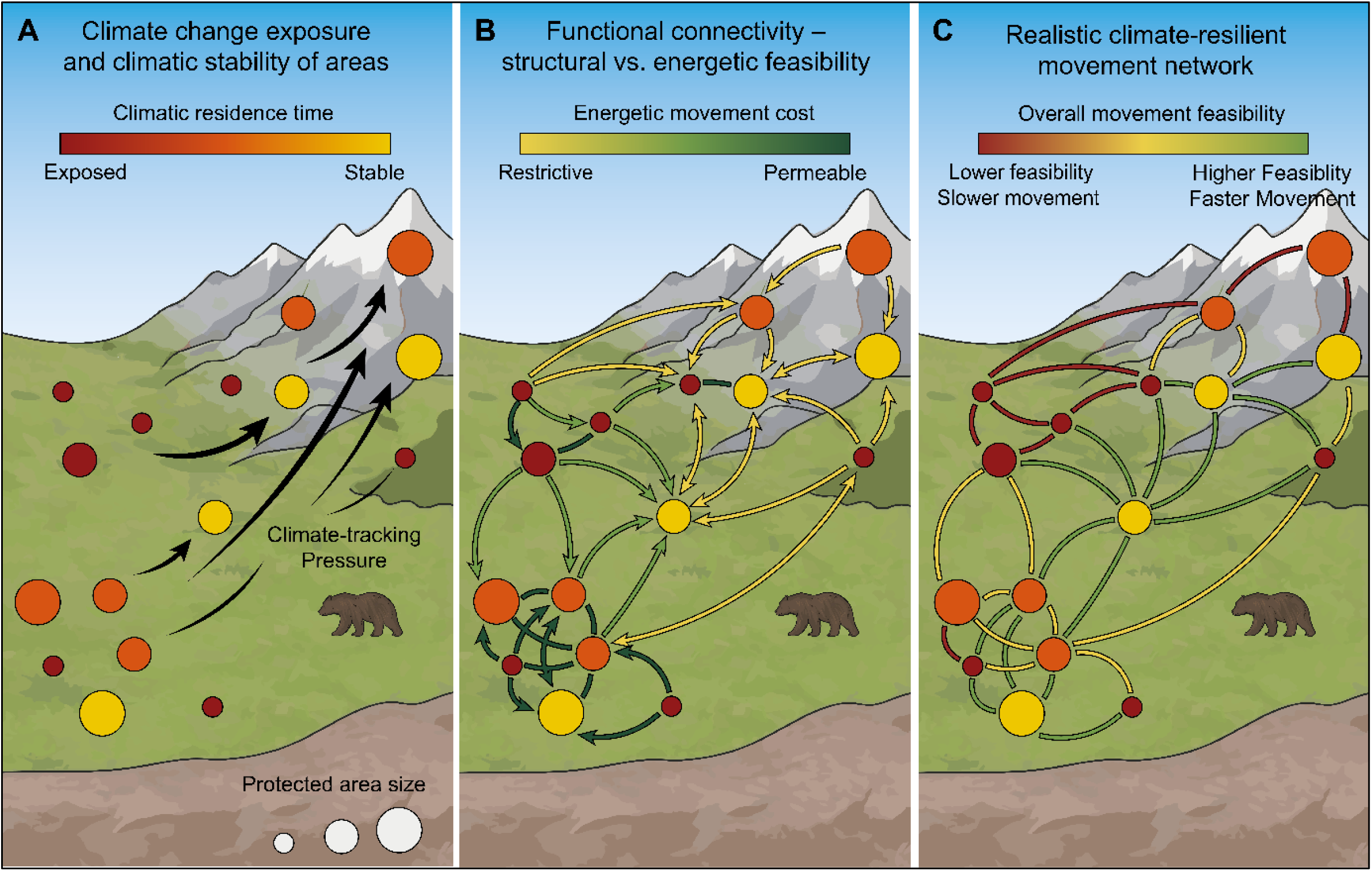
Climate-resilient connectivity framework for protected areas. (A) Protected areas (PAs) vary in climatic residence time, ranging from climate-exposed sites (red) to more climatically stable or refugial sites (yellow–green). Circle size represents PA area, illustrating that smaller PAs may provide less internal climatic buffering than larger ones and may therefore become unsuitable more quickly under climate change. These spatial differences create directional pressure for climate tracking toward more stable areas. (B) Functional connectivity is shaped by energetic movement costs across the landscape. The gradient from permeable (low-cost) to restrictive (high-cost) conditions shows how terrain and land use influence movement feasibility, emphasizing that structural connectivity alone may overestimate realistic dispersal pathways. (C) Integrating climatic stability, PA size, and energetic resistance identifies a climate-resilient movement network, highlighting where movement among climate-exposed and climatically stable PAs is more or less feasible.

## Results

### Geographic Distribution of Climate-Exposed and Climate-Resilient PAs

Climate velocity revealed strong spatial heterogeneity in climatic residence time across Europe’s protected-area network (Fig. 2). Protected areas with comparatively short residence times were most prominent across broad parts of western, eastern, and northern Europe, with many occurring in relatively low-relief landscapes where climatic conditions are projected to shift rapidly. This pattern was especially evident across extensive lowland areas, including parts of the western European lowlands and Central European plains, as well as more easterly and northerly regions shown in Fig. 2. These areas likely reflect the combined effects of weak spatial climatic gradients and rapid warming, which increase horizontal climate velocity and reduce the potential for long-term in situ persistence despite formal protection. In contrast, protected areas with longer residence times were concentrated in major mountain systems, where steep elevational gradients are associated with greater climatic stability (Fig. 2). High-resilience PAs were consistently associated with the Alps, Pyrenees, Carpathians, Cantabrian Mountains, Apennines, and the Scandinavian highlands. Strong elevational and topographic gradients in these regions reduce effective climate velocity by enabling relatively short-distance tracking of suitable conditions, increasing the likelihood that mountainous PAs function as long-term climatic refugia under projected warming trajectories.

**Figure 2.**
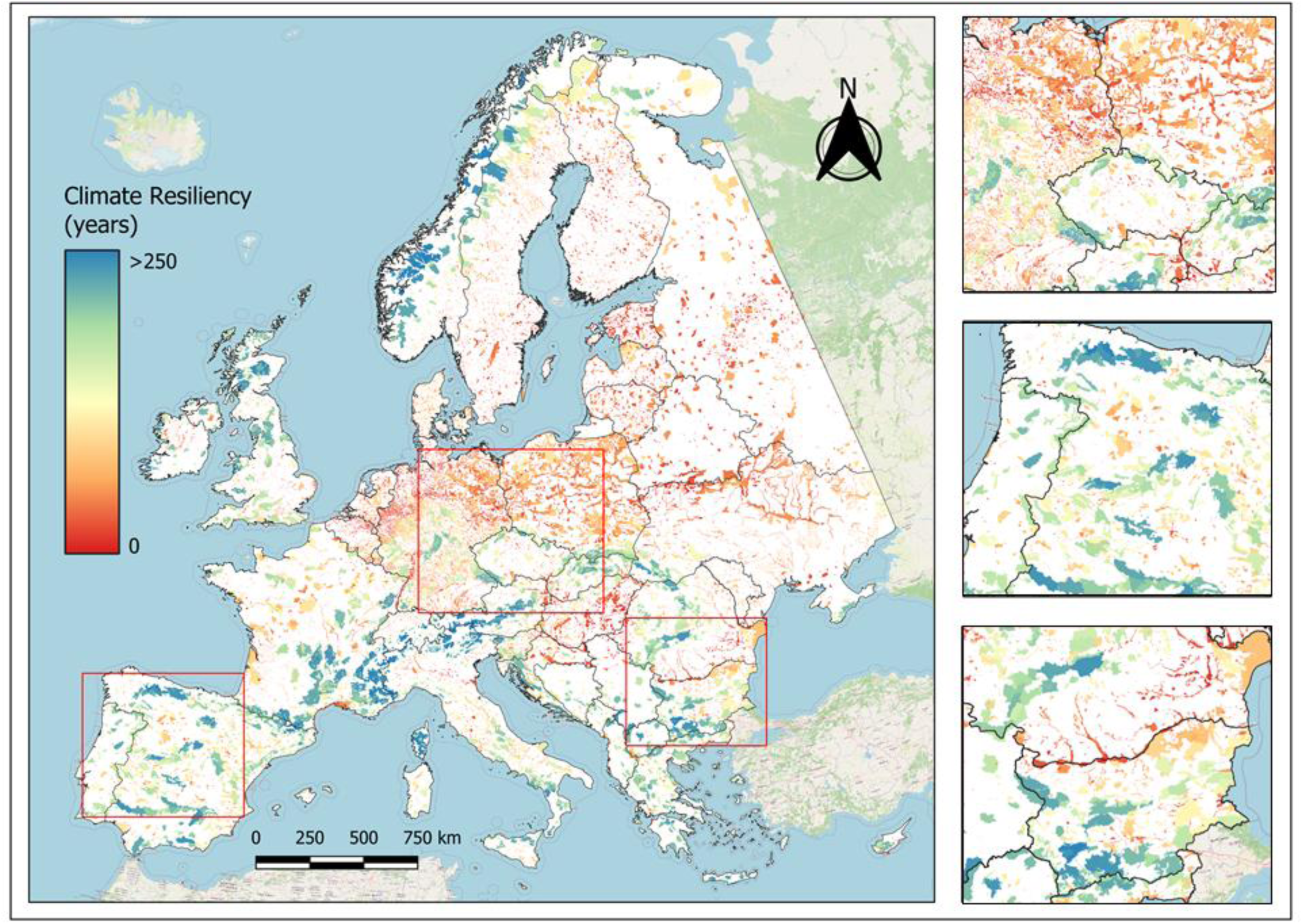
Spatial distribution of climatic resilience across Europe’s terrestrial protected areas. Climatic resilience is measured as climatic residence time (years), indicating the projected duration over which current climatic conditions persist within each protected area. Colours range from low resilience (red) to high resilience (blue). Insets show enlarged views of the outlined regions and illustrate strong spatial heterogeneity across Europe. In general, lowland protected areas exhibit shorter residence times, whereas mountainous and topographically heterogeneous areas show greater climatic stability and stronger potential to act as climatic refugia. Larger protected areas also tend to have longer residence times, reflecting greater internal climatic buffering capacity.

### Spatial Patterns of Energy-Based Functional Connectivity Among Herbivores and Carnivores

We next evaluated how movement energetics and habitat suitability jointly structure functional connectivity across Europe for generalized herbivore and carnivore archetypes (Fig. 3). Current density revealed strong spatial organization and marked differences between archetypes. For herbivores, connectivity was comparatively broad and continuous across much of Europe, with relatively high movement potential extending across Fennoscandia, parts of Iberia, and several mountain and foothill systems in central, southern, and eastern Europe (Fig. 3). Lower current density was concentrated in highly urbanized areas and across extensive intensively managed lowlands, where movement pathways became more diffuse or weakly connected. In contrast, carnivore connectivity was much more restricted and spatially channeled (Fig. 3). High current density was concentrated mainly in northern Europe and along major mountain and foothill belts (especially in southern Europe). In contrast, large parts of western and central Europe were characterized by persistently low movement potential. This fragmentation was especially pronounced across the British Isles and broad lowland regions spanning northern France, Benelux, Germany, and adjacent agricultural plains. Across both archetypes, topographically complex and less intensively modified landscapes supported the strongest connectivity, while flat, human-dominated lowlands acted as major constraints on movement, with these limitations substantially stronger for carnivores.

**Figure 3.**
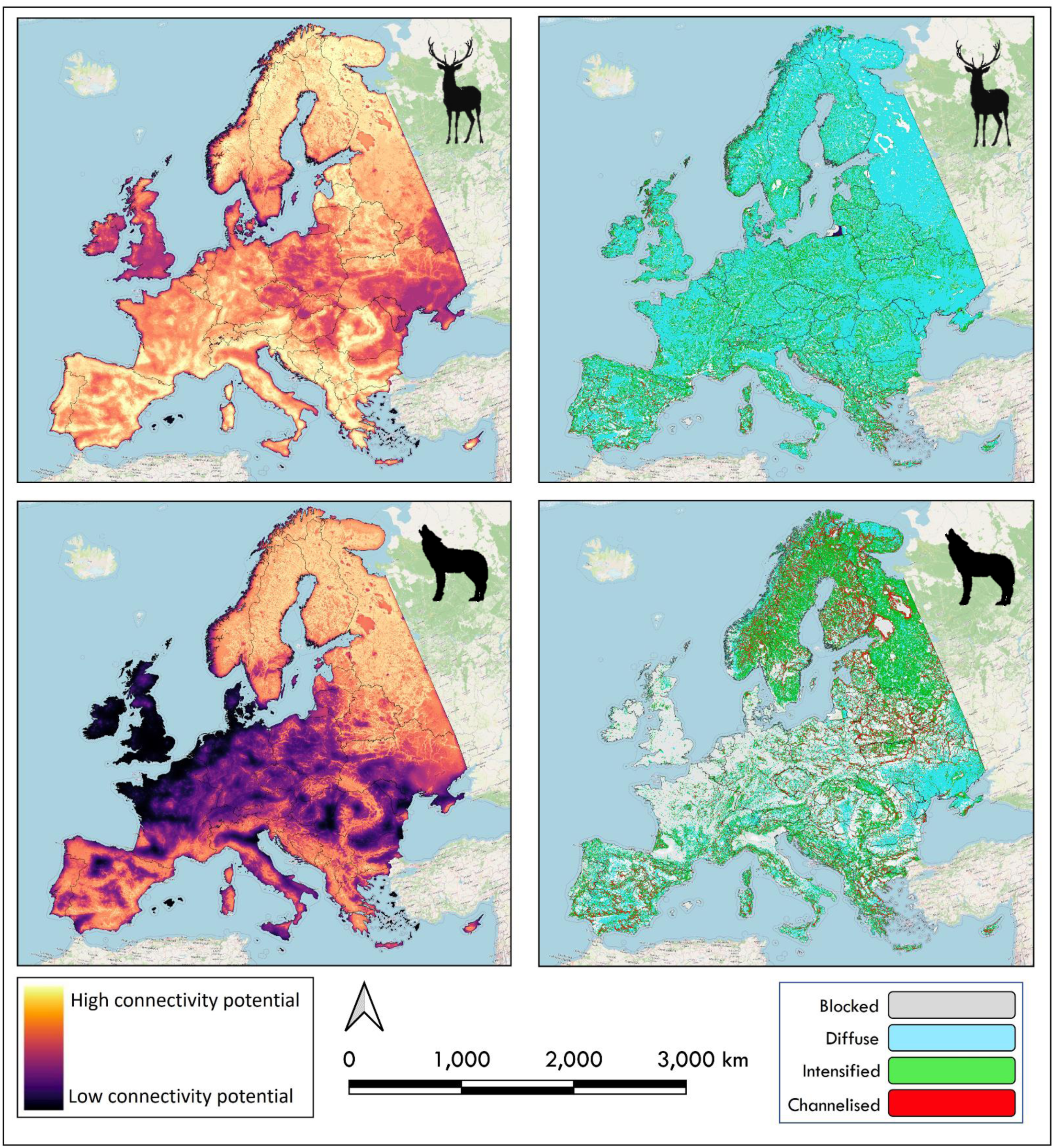
Functional connectivity patterns across Europe for generalized herbivore and carnivore archetypes. Top-row panels represent the herbivore archetype and bottom-row panels the carnivore archetype. Left panels show continuous Omniscape current density, with higher values indicating greater connectivity potential and lower movement resistance. The right panels show normalized connectivity classes identifying blocked, diffuse, intensified, and channelized movement areas. Connectivity estimates combine habitat-suitability–based permeability with energetic movement costs and resistance from human-modified land uses. Compared with herbivores, carnivores show more spatially restricted and channelized connectivity, whereas herbivore connectivity is broader and more diffuse across Europe.

### Continental Patterns in the Co-occurrence of Climatic Resilience and Functional Connectivity

Overlaying energy-based functional connectivity with PA climatic resilience revealed a clear geographic structure in the co-occurrence of movement potential and climatic stability across Europe (Fig. 4). For both archetypes, the most favourable combinations of high connectivity and high climatic resilience were concentrated in topographically complex, predominantly mountainous regions, whereas extensive lowland areas were dominated by low connectivity and low resilience, indicating places where rapid climatic turnover coincides with strong movement constraints (Fig. 4). In addition to this broad mountain–lowland contrast, we show that favourable overlap is organized along major mountain arcs rather than uniformly across Europe (Fig. 4). For herbivores, high-connectivity, high-resilience combinations recur along the Scandinavian mountains, the Pyrenees–Cantabrian system, the Alps, the Apennines, and especially the Dinaric–Balkan region, with further patches in the Carpathians (Fig. 4). By contrast, the western and central European lowlands are dominated by unfavourable combinations, with only isolated favourable pockets. These patterns are also strongly heterogeneous within regions: even in mountain systems, climatically resilient connectivity is patchy and interspersed with intermediate and unfavourable areas. This patchiness is accentuated for carnivores, for which favourable overlap is more restricted and fragmented, and many areas that appear mixed to favourable for herbivores shift toward lower-connectivity combinations (Fig. 4). Together, these patterns indicate that climate-tracking opportunities are concentrated in a limited set of mountainous corridors and transitional foothill landscapes, whereas much of lowland Europe remains both climatically exposed and functionally difficult to traverse.

**Figure 4.**
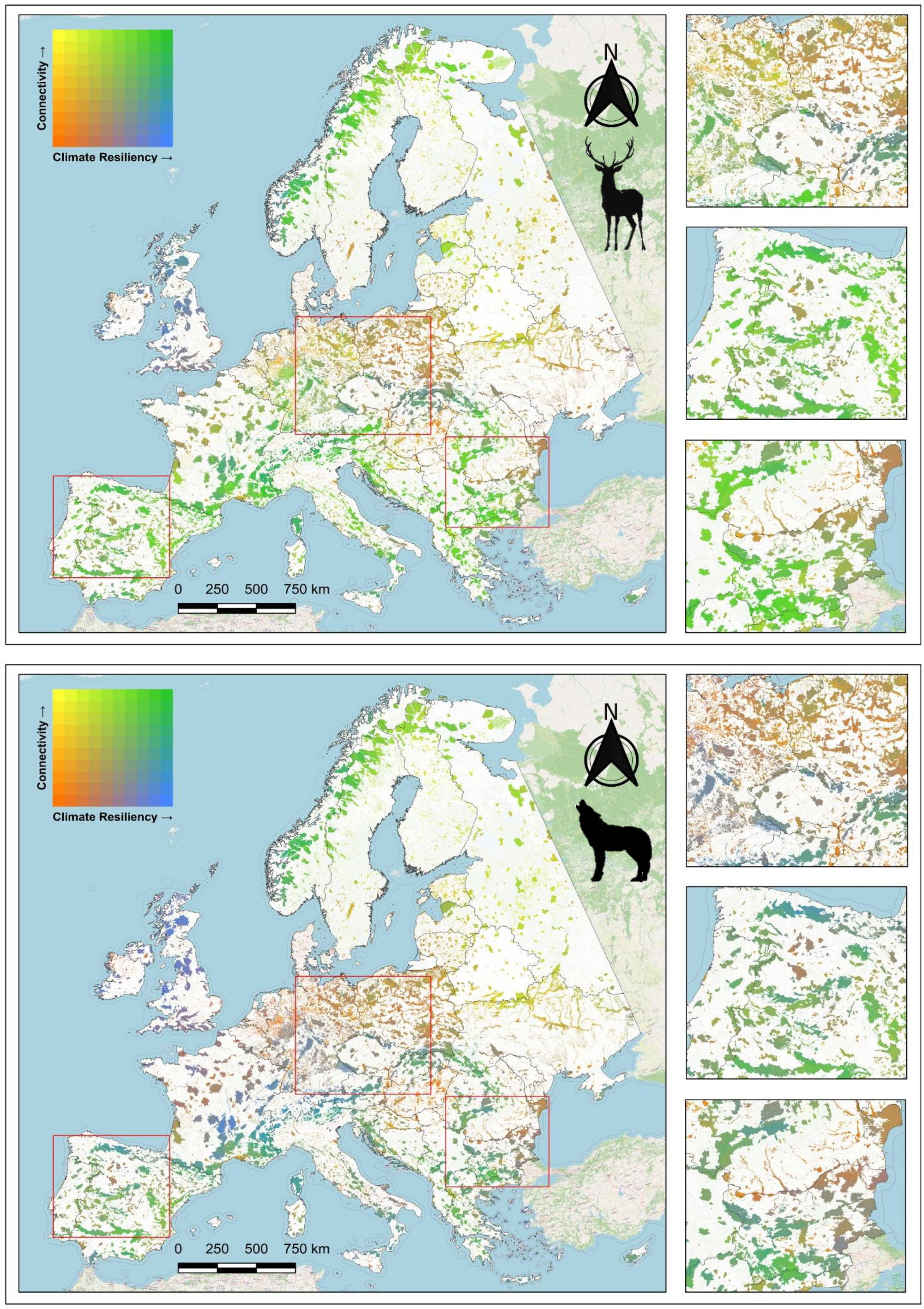
Climatic resilience and energy-based functional connectivity across European protected areas. Spatial analysis shows the overlap between current energy resistance density–based connectivity and protected-area climatic resilience, measured as climatic residence time. Colours indicate bivariate combinations of connectivity and climatic resilience: green, high connectivity and high resilience; orange, low connectivity and low resilience; blue, high resilience but low connectivity; and yellow, high connectivity but low resilience. The top panel represents the generalized herbivore archetype and the bottom panel the generalized carnivore archetype. Insets show enlarged views of the outlined regions and highlight spatial heterogeneity in the co-occurrence of movement potential and climatic stability across Europe.

This contrast was also reflected in the country-level summaries (Fig. 5) and the within-country PA distributions (Fig. S1), which show substantial heterogeneity and frequent mismatches between the two dimensions rather than a simple linear relationship. For herbivores, this alignment was most evident in southern and mountainous countries such as Montenegro, Albania, Bosnia and Herzegovina, and North Macedonia, which occupy the high-connectivity, high-resilience end (Fig. 5). In contrast, the United Kingdom, Ireland, Denmark, and the Netherlands cluster toward the low-connectivity, low-resilience end, while Latvia, Estonia, Finland, and Lithuania illustrate a recurrent mismatch in which relatively high connectivity coincides with comparatively low climatic resilience (Fig. 5). For carnivores, the same broad pattern persists, but connectivity is more spatially restricted, with Montenegro, Bosnia and Herzegovina, Albania, and Austria standing out as relatively connected and stable, whereas the United Kingdom, Ireland, Belgium, and Germany remain low in connectivity and resilience (Fig. 5). Northern and eastern countries such as Finland, Estonia, Latvia, and Russia again show comparatively high connectivity despite lower climatic resilience, while Portugal, Spain, and Switzerland occupy intermediate to relatively high positions on both axes (Fig. 5). Together, these results show that climatic stability and connectivity align most strongly in mountainous southern Europe, whereas many western, central, northern, and eastern countries are limited by low movement potential, high climate exposure, or both.

**Figure 5.**
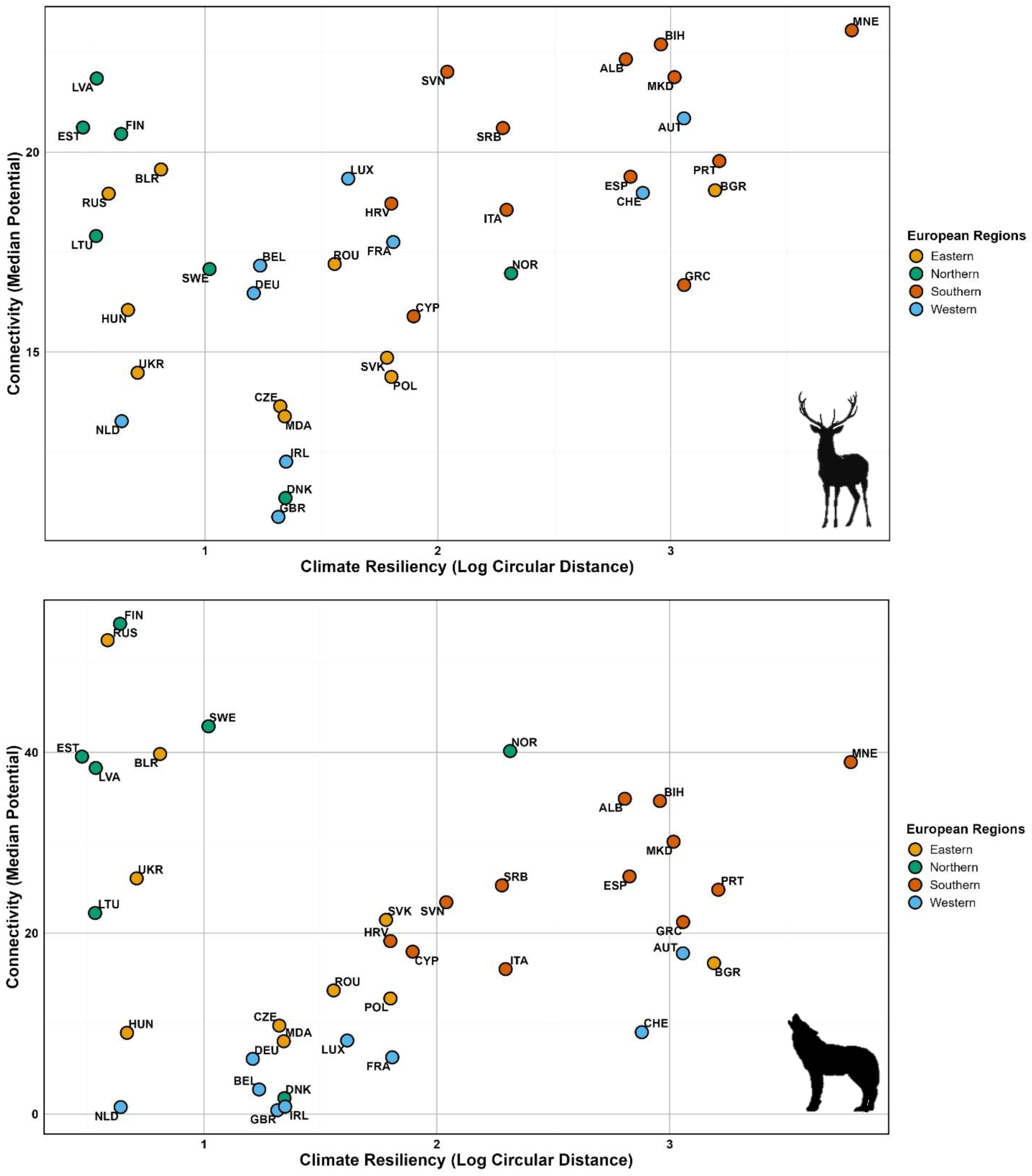
Country-level variation in climatic resilience and energy-based functional connectivity across Europe. Scatterplots show country-level connectivity, measured as median potential, against climatic resilience, expressed as the log circular climate displacement distance, for the generalized herbivore archetype (top panel) and generalized carnivore archetype (bottom panel). Each point represents a country and is labeled with its ISO 3166-1 alpha-3 code. Points are coloured by broad European region (Eastern, Northern, Southern, and Western). The plots highlight substantial cross-national heterogeneity and contrasting connectivity distributions between herbivore and carnivore archetypes.

## Discussion

### Refugia accessibility and climate-resilient connectivity

Our study shows that a substantial fraction of European protected areas (PAs) is likely to lose the climatic conditions for which they were originally designated in the coming decades, while many are also insufficiently connected to support species redistribution under climate change. This reinforces and extends continental and global assessments showing accelerating climate exposure and declining climatic residence times across protected-area networks (Araújo et al., 2011; Cimatti et al., 2025; Elsen et al., 2020; Heller & Zavaleta, 2009; Parks et al., 2022). Similar patterns have already been reported for Alpine systems, the Iberian Peninsula, and Central Europe, where climatic analogues for many PAs are increasingly projected to occur beyond their present boundaries (Mingarro & Lobo, 2021; Semenchuk et al., 2021). However, identifying exposed PAs and potential refugia is only part of the challenge. The long-term persistence of many PAs, particularly in lowland Europe, will depend on whether species can actually move through functionally connected landscapes toward areas where suitable climates are expected to persist (Ranius et al., 2023). By integrating climate metrics with movement energetics, our framework (Fig. 1) helps bridge this gap. It clarifies not only where climate-driven movement is likely to become necessary, but also where such movement is more likely to be feasible at broad spatial scales (Fig. 4). In doing so, it moves beyond the simple identification of refugia and provides a basis for more actionable climate-resilient connectivity planning (Buenafe et al., 2025; Carroll et al., 2018; Keeley et al., 2018).

In this context, mountain systems are particularly important for climate adaptation. Steep spatial climatic gradients in these regions can create shorter and more attainable climate-tracking pathways, thereby increasing climatic residence times and strengthening their potential role as refugia (Cimatti et al., 2025; Dragonetti et al., 2024; Heikkinen et al., 2020). Yet this conservation value is conditional on accessibility. Climatically stable mountain PAs cannot serve as effective endpoints for climate tracking if surrounding landscapes do not permit movement into them (Morelli et al., 2016, 2017). As a result, the adaptive value of mountain refugia depends not only on local climatic stability, but also on the permeability and configuration of the broader landscape matrix (Zeller et al., 2020). This makes foothill and transitional landscapes especially important because they often serve as links between climatically exposed lowlands and more stable montane regions (Caballero-Díaz et al., 2025; Pan et al., 2025). Where these connecting landscapes are fragmented or highly resistant to movement, even well-positioned refugia may remain functionally out of reach (Caballero-Díaz et al., 2026; Moniem et al., 2025). In that sense, the effectiveness of climate adaptation depends not only on where refugia occur, but also on whether protected areas are large enough, well enough connected, and embedded within landscapes that allow redistribution to unfold in practice (Anderson et al., 2023; Costanza & Terando, 2019; McGuire et al., 2016).

### The limits of static protection: fragmentation and the feasibility of climate-tracking movement

A complementary vulnerability emerges where rapid climatic turnover coincides with the small size and fragmented configuration of many European protected areas (Fig. 2). As mentioned, mountain refugia can support climate adaptation only if they are reachable; for many lowland PAs, this is highly impractical (Fig. 4). Small PAs provide limited internal climatic buffering because their restricted area constrains how far species can track shifting conditions before reaching site boundaries, increasing reliance on dispersal over relatively short timescales (Geyer et al., 2017; Ranius et al., 2023; Rannow & Neubert, 2014). This problem is especially acute in Europe, where much of the PA network, particularly many Natura 2000 sites, consists of small, spatially discontinuous units embedded in intensively used landscapes (Evans, 2012; Mazaris et al., 2013).

Under climate change, the key issue is therefore no longer simply whether biodiversity is best conserved in one large reserve or several small ones, but whether species can move among sites as climates shift (Hilty et al., 2020; Keeley et al., 2018; Littlefield et al., 2019). Accordingly, the effectiveness of Europe’s PA network depends not only on the total protected area, but also on patch size, topographic context, and network configuration (Doxa et al., 2022; Hlásny et al., 2021). Expanding or clustering nearby sites may modestly increase climatic heterogeneity and extend residence times, but this will often be insufficient where climatic analogues lie beyond reachable distances (Parks et al., 2023). In such settings, persistence increasingly depends on whether climate-exposed PAs are embedded within landscapes that allow movement toward more stable areas (Hilty et al., 2020; Sonntag & Fourcade, 2022). Without barrier removal, permeability restoration, and functional links among sites, many small and fragmented PAs in intensively modified landscapes may remain protected in formal terms while becoming climatically obsolete in practice.

This is precisely why the vulnerability of small, fragmented PAs cannot be understood solely in terms of structural connectivity (Fig. 1). A central contribution of our study is to show that landscapes that appear continuous or permeable are not necessarily traversable in practice, because movement feasibility depends not only on land-cover structure but also on the energetic costs of moving across it (Araújo et al., 2025). Incorporating movement energetics, therefore, reveals a more constrained picture of climate-tracking potential: flat, intensively modified lowlands may require long-distance movement to track suitable climates, while topographically complex terrain can also impose substantial energetic costs over short distances (Berti et al., 2025; Wilson et al., 2012). Accordingly, a feasible climate-tracking movement is concentrated where climatic gradients, landscape permeability, and manageable energetic costs align, especially in foothill and transitional settings that link exposed lowlands with more stable uplands (Fig. 4; Cordes et al., 2025). This highlights energetic resistance as a broad ecological filter on climate adaptation and suggests that restoration should focus less on maximizing continuity everywhere than on improving movement feasibility in the parts of the landscape that matter most, for example through stepping stones, barrier mitigation, and more permeable land-use mosaics (Cameron et al., 2022; Costanza & Terando, 2019; McRae et al., 2012). Framed this way, the value of connectivity planning lies not simply in linking habitats but in identifying where targeted restoration and cross-border coordination are most likely to sustain climate-tracking movement under ongoing change, an issue that is directly relevant to current European conservation policy and implementation.

### Policy implications, study limitations, and future directions

The spatial patterns revealed by our analysis address a key implementation gap in European biodiversity policy. Although the EU Biodiversity Strategy for 2030, the Trans-European Nature Network (TEN-N), and the EU Nature Restoration Regulation (NRR) all emphasize the need for a coherent and well-connected ecological network, they provide limited guidance on where connectivity is most critical under accelerating climate change (Bores et al., 2024; Liquete et al., 2024; Regulation (EU) 2024/199). Our analyses suggest that conservation planning should shift from prioritizing where habitat appears structurally continuous to prioritizing where movement remains climatically and energetically feasible under ongoing change. This is especially important in Europe, where many ecologically relevant movement pathways cross national borders, while governance and implementation remain fragmented (Leibenath et al., 2010; Rüter et al., 2014; Sluis, 2022). Where the persistence of climate-exposed PAs depends on access to more stable parts of the landscape, embedding energy-informed, climate-resilient connectivity into restoration prioritization, transboundary corridor strategies, and national implementation frameworks could better direct investments toward the parts of the network most likely to sustain climate-tracking movements. Existing initiatives such as the European Green Belt show that such cross-border coordination is both feasible and effective when conservation goals are aligned across jurisdictions (Zmelik et al., 2011). In this sense, the main policy implication of our study is that climate adaptation depends less on maximizing continuity everywhere than on strategically reinforcing the regional conditions most likely to sustain feasible movement over time (Bauduin et al., 2020; Carter et al., 2024; Schloss et al., 2022; Stark et al., 2025).

At the same time, our framework should be understood as a continental screening tool rather than a predictor of realized dispersal. Its strength lies in identifying where climate exposure, energetic constraints, and fragmented protection are most likely to interact, but this generality also imposes clear limitations. Our analyses rely on generalized movement–energetics models that do not capture species-specific dispersal traits, behaviours, or physiological tolerances (Berti et al., 2022), and our resistance surfaces do not incorporate behavioural avoidance, mortality risk, demographic feedbacks, or fine-scale microclimatic heterogeneity that may alter movement outcomes. We also did not assess whether recipient refugia will retain sufficient habitat quality or carrying capacity to support incoming populations over time (Gillingham et al., 2024; Sinclair et al., 2025), and our broad-scale representation of landscapes may overlook local restoration features, fine-scale barriers, or ongoing interventions that could facilitate or constrain redistribution (Ellis et al., 2025; Rao et al., 2025). These caveats do not weaken the central message of the study; rather, they define its intended scope and point directly to future research needs. Building on this foundation will require integrating species-specific dispersal and demographic processes to translate coarse-filter constraints into taxon-specific risk assessments (Landim et al., 2025; Reyes-Moya et al., 2022), as well as coupling energetic connectivity with land-use change and alternative restoration pathways under different socio-economic futures (Araújo et al., 2025). In conclusion, we suggest that restoration that ignores movement feasibility risks creating habitat that is suitable but functionally unreachable, whereas integrating climate dynamics with energetic constraints can support more adaptive and forward-looking network design.

## Methods

### Climate Velocity and Resilience of Protected Areas

We quantified the rate, direction, and spatial heterogeneity of climate change across Europe to identify protected areas (PAs) exposed to rapid climatic turnover versus those that are comparatively resilient (Fig. 1A). Climate change velocity was calculated using a gradient-based approach that relates the temporal rate of temperature change to local spatial climatic gradients (García Molinos et al., 2019; Loarie et al., 2009). Mean annual near-surface air temperature data were obtained from CHELSA v2.1 at 30-arc-second (∼1 km²) resolution for a historical baseline period (1981–2010) and for future projections (2041–2070) under the SSP3–7.0 scenario, based on three Earth system models (GFDL-ESM4, UKESM1-0-LL, and MPI-ESM1-2-HR; Karger et al., 2023; Karger & Zimmermann, 2021). We selected SSP3–7.0 since it represents a policy-relevant high emission trajectory suitable for identifying near-term gradients of climatic exposure (Meinshausen et al., 2020; Popp et al., 2017). Temporal temperature gradients were computed as the annualized difference between baseline and future mean temperatures, and spatial gradients were estimated using a 3 × 3 cell neighborhood and Horn’s finite-difference method (Horn, 1981).

Climate velocity was calculated as the ratio of temporal to spatial gradients, yielding an estimate of the horizontal displacement (km yr⁻¹) of climatic conditions (for more details, see supplementary materials). These velocity estimates form the basis for quantifying PA-scale climatic stability and are subsequently used to derive metrics of climatic resilience and exposure, including climate residence time. To account for uncertainty in climate projections, velocity estimates were averaged across three Earth system models: GFDL-ESM4, UKESM1-0-LL, and MPI-ESM1-2-HR (Cimatti et al., 2025). Climate velocity values were then spatially intersected with terrestrial protected areas from the World Database on Protected Areas, excluding marine and coastal sites (Bingham et al., 2019). Protected-area climatic resilience was quantified using climate residence time, defined as the expected duration over which a PA retains its current climatic conditions (Brito-Morales et al., 2018). Residence time was estimated by dividing the effective radius of each PA (assuming a circular geometry) by its mean internal climate velocity (following methodology from Loarie et al., 2009). Longer residence times indicate greater climatic stability and potential refugial function, whereas shorter times indicate rapid climatic turnover and increased reliance on dispersal to maintain population (Haight & Hammill, 2020).

### Energetic Cost of Terrestrial Movement and Landscape Resistance

We constructed landscape resistance layers to evaluate whether climate-driven redistribution among protected areas is both physiologically and structurally feasible (Fig. 1B). To quantify the energetic constraints on movement, we first estimated the cost of terrestrial locomotion across Europe using a biophysical movement–energetics framework based on the Enerscape formulation, which links locomotion energy expenditure to terrain slope via an allometric cost-of-transport relationship (Fig. S2; Berti et al., 2022). Because the full formulation includes body mass, we applied a normalized variant in which energetic costs depend only on slope, enabling inference across a broad range of terrestrial mammals (Berti et al., 2022). Concretely, we computed a normalized energetic cost function g(θ), where θ is the terrain slope (degrees). This normalization rescales energetic expenditure between theoretical minimum and maximum values to yield a unitless index of relative movement cost that is independent of body mass. The resulting “energy landscape” therefore captures the spatial pattern of energetic difficulty imposed by terrain itself rather than species-specific metabolic rates (Fig. S1). We derived the slope at 1 km resolution from EU-DEM elevation data and calculated g(θ) for each grid cell.

To represent landscape permeability and habitat suitability, we developed continental-scale species distribution models (SDMs) for a set of focal mammals with transboundary European ranges and contrasting ecological requirements (Fig. S1). We modelled four predators, brown bear (*Ursus arctos*), wolf (*Canis lupus*), Eurasian lynx (*Lynx lynx*), and wolverine (*Gulo gulo*), and six herbivores/species groups, elk (*Alces alces*), red deer (*Cervus elaphus*), wild boar (*Sus scrofa*), caribou (*Rangifer tarandus*), chamois (*Rupicapra* spp.), and ibex (*Capra* spp.), with mountain ungulates grouped at the genus level. Using presence-only data, we fit generalized linear mixed models with covariates describing proportional land cover, elevation, and climatic conditions to estimate habitat suitability for each taxon. We then performed model averaging across all top-supported models (ΔAICc < 8; Burnham & Anderson, 2002), inverted the resulting suitability scores, and transformed them into resistance layers so that high suitability corresponded to low resistance, and vice versa. Because SDMs may not fully capture discrete, high-resistance features that strongly impede movement (e.g., large water bodies or engineered barriers), we increased realism by “burning in” additional resistance for major rivers and lakes, glaciers, motorways, and border barriers (see Supplementary Materials for full details).

Finally, we harmonized the slope-derived energetics surface and the SDM-derived resistance surface by scaling each to a common range (1–1000) and combining them pixel-wise using the maximum of the two values to generate a composite resistance layer (Fig. S1). This composite surface represents the relative resistance a moving animal would experience by jointly accounting for energetic expenditure, land use, and human infrastructure. In doing so, it differentiates landscapes where movement is energetically efficient from areas where dispersal would impose high physiological demands even in the absence of explicit barriers. Overall, the resulting resistance layer provides an indicator of energetic resilience, highlighting regions most capable of supporting sustained movement under ongoing climate change.

### Connectivity Analysis Integrating Energy and Landscape Structure

We integrated energetic resistance into the connectivity analysis by modelling landscape connectivity with Omniscape, an omnidirectional circuit-theory approach that uses a moving-window framework to estimate connectivity potential continuously across the landscape (Landau et al., 2021), rather than relying on predefined source–destination nodes such as protected areas (Fig. 1B). In circuit theory, the resistance surface is represented as an electrical network, where current flow reflects the probability of movement distributed across all plausible pathways, not a single least-cost route (McRae et al., 2008). The resulting cumulative current-density surfaces highlight areas of relatively high and low movement potential and identify where connectivity is most limited (Dickson et al., 2019). To summarize connectivity across taxa, we combined cumulative current density layers into two archetypal species groups (predators and herbivores) by taking the maximum current value per pixel within each group, yielding continent-wide, geographically explicit representations of movement potential. We then quantified connectivity in and around each protected area by generating 25 km buffers around all WDPA-derived polygons (n = 10,908) and extracting mean and median cumulative current density within each buffer using zonal statistics for both species groups (see Fig. S3 in supplementary materials). Using buffers extending beyond protected area boundaries was essential to capture the surrounding landscape context and the potential for dispersal through and beyond individual protected areas (Fig. 1C).

## Acknowledgments

We are grateful to Marta Cimatti for insightful discussions during this study’s design.

## Authors’ contributions

Conceptualization: G.S, N.F, H.M.P; Formal analysis: JD, NRP, EB; Investigation: JD, NRP, EB; Methodology: all authors; Project administration: GS & H.M.P; Supervision: N.F and H.M.P; Visualization: JD, NRP, ACMG, GS; Writing – original draft: G.S; Writing – review & editing: all authors.

## Data and Code Availability

All data and code used in this study are available on GitHub (https://github.com/GavinStark89/VECTOR-Velocity-based-Energetics-informed-Connectivity-for-Tracking-Organisms-to-Refugia) and archived on Zenodo (DOI: 10.5281/zenodo.19609128).

