## Supplementary material for "The Hidden Costs of Climate Change Tracking: Climate Velocity, Movement Energetics, and Connectivity in European Protected Areas": SM

### Protected-area dataset, buffering, and regional summaries

Protected-area boundaries were obtained from the World Database on Protected Areas (WDPA; [July 2025 release, accessed on 12 January 2026]) and filtered to retain only terrestrial sites, excluding marine and coastal areas, yielding a final analytical dataset of 10,908 polygons. These polygons were used in two complementary ways: first, the original protected-area boundaries were used to extract mean internal climate velocity and estimate climatic residence time; second, 25 km buffers around each polygon were used to extract mean and median cumulative current density for both archetypal groups using zonal statistics, thereby capturing the surrounding landscape matrix through which climate-tracking movement into and out of protected areas would occur. These protected-area level summaries formed the basis for comparative analyses of climatic resilience and functional connectivity across Europe, including country-level summaries visualized in the main manuscript using ISO 3166-1 alpha-3 country codes and broad regional groupings (Fig. S1).

**Figure S1**. Country-level relationships between climatic resilience and energy-based functional connectivity. Panels show scatter plots of mean current density (energy-based functional connectivity) versus climatic resilience, expressed as the log of circular climate displacement distance, for individual European countries (ISO 3166-1 alpha-3 codes). Each point represents a protected area within a country, illustrating within-country variation in the association between climatic resilience and functional connectivity. The distributions highlight substantial heterogeneity both within and among countries in the co-occurrence of climatic stability and connectivity across Europe.

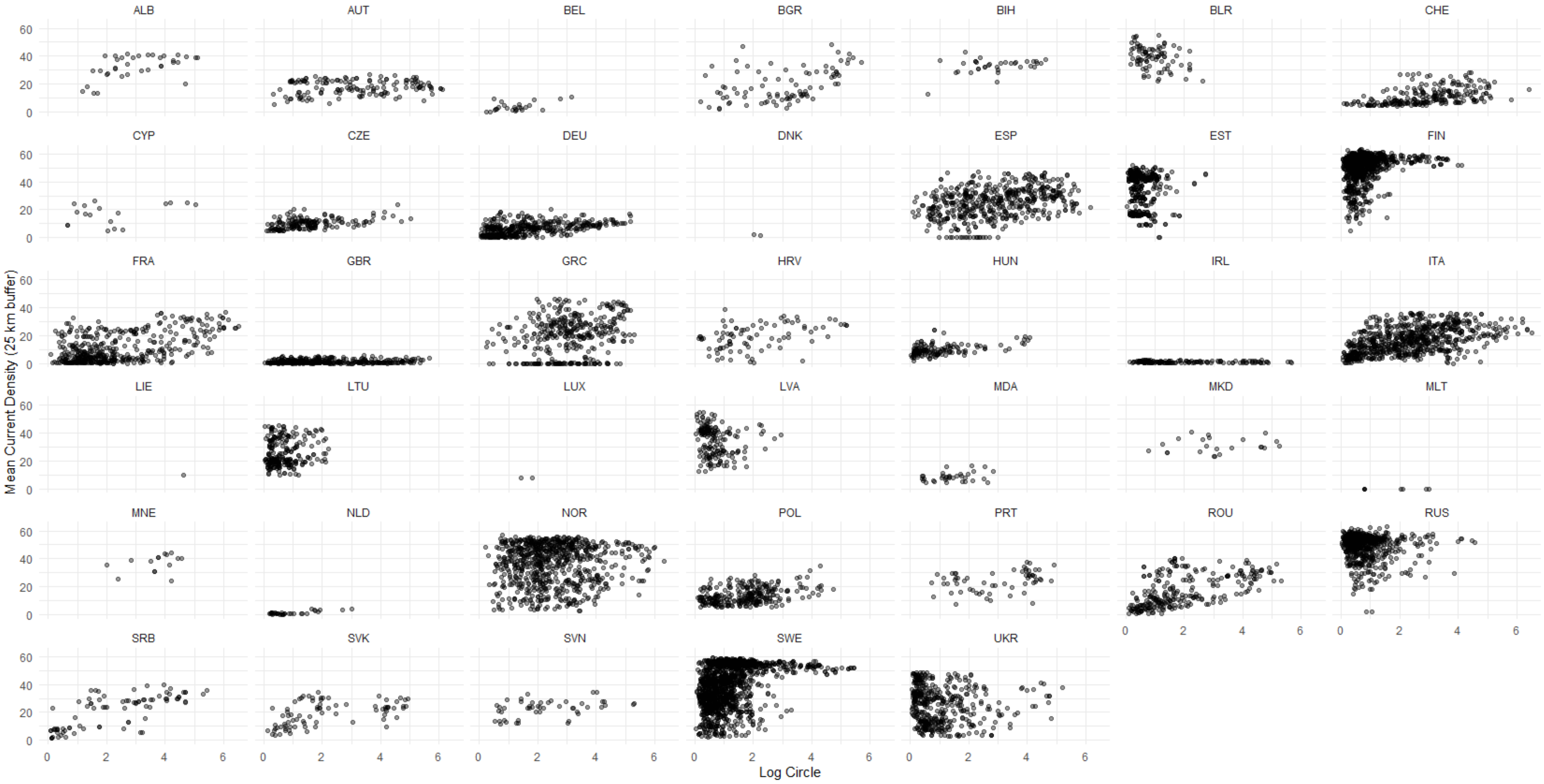

### Climate-velocity estimation calculation

Local climate velocity was then calculated as the ratio of temporal to spatial gradients, yielding an estimate of the horizontal displacement of climatic conditions in km yr⁻¹. This metric describes the annual distance species would need to move to maintain constant climatic conditions under projected warming:

$$Local\ Velocity\ = \frac{slope\ \left(\frac{unit}{year}\right)}{spatial\ gradient\ \left(\frac{unit}{km}\right)} = gVoCC\ (km\ yr^{-1})$$

For a specific variable (e.g., temperature in °C), the equation can be adapted; and would typically be written as:

Local Velocity= $\frac{\mathbf{\partial T/\partial t}}{\Delta t}$

where $\partial T/\partial t$ is the temporal trend (°C $yr^{-1}$) and the delta T is the local spatial gradient (°C $km^{-1}$).

Temporal temperature gradients were computed as the annualized difference between baseline and future mean temperature, and spatial temperature gradients were estimated using a 3 × 3 moving window and Horn's finite-difference method.

$$G_t = \frac{(Tfuture - Thistorical)}{\Delta t}$$

where *T(future)* and *T(historical)* represent mean annual temperatures for the future and baseline periods, respectively, and $\Delta t$ is the number of years between their midpoints. The spatial temperature gradient ($G_s$, °C· $km^{-1}$) was computed using Horn's method for surface slope:

$$G_S = \sqrt{\left(\frac{\partial T}{\partial x}\right)^2 + \left(\frac{\partial T}{\partial y}\right)^2}$$

where $\partial T/\partial x$ and $\partial T/\partial y$ are longitudinal and latitudinal temperature gradients calculated within a 3 × 3 cell neighborhood. Local climate velocity ($Vc$, km·$yr^{-1}$) was then derived as the ratio between the temporal and spatial gradients:

$$Vc = \frac{Gt}{Gs}$$

This represents the instantaneous horizontal velocity of temperature change, indicating the distance that species or ecosystems must move annually to maintain constant climatic conditions. Calculations were implemented in R using the VoCC package (García Molinos et al., 2019), and ensemble means across three Earth System Models were used to account for structural and parametric uncertainty.

### Protected-area climatic resilience and residence time

Protected-area climatic resilience was quantified using climatic residence time, defined as the expected duration over which a protected area retains its current climatic conditions. For each terrestrial protected area (from the World Database on Protected Areas; UNEP-WCMC & IUCN, 2023), we extracted the mean internal climate velocity (*Vc*) and estimated the effective radius assuming a circular geometry. Residence time ($R_t$, years) was then calculated by dividing the protected-area effective radius by the mean internal climate velocity. Higher values, therefore, indicate greater climatic stability and greater potential for refugial function, whereas lower values indicate more rapid climatic turnover and greater reliance on redistribution to maintain populations. We estimated climate residence time as:

$$Rt = \frac{r}{Vc}$$

where *r* is the PA's radius (km), when forcing a circular shape. This metric reflects the expected time before a PA's climatic conditions are fully displaced, integrating both spatial extent and climate velocity. Marine and coastal PAs were excluded to focus on terrestrial systems. The resulting dataset provides a spatially explicit estimate of climate velocity and protected area resilience across Europe.

### Focal taxa, occurrence records, and taxonomic grouping

To represent broad differences in movement ecology and landscape use, we modelled a set of large terrestrial mammals with transboundary European distributions and contrasting ecological requirements. The carnivore group comprised brown bear (*Ursus arctos*), wolf (*Canis lupus*), Eurasian lynx (*Lynx lynx*), and wolverine (*Gulo gulo*). The herbivore group comprised elk (*Alces alces*), red deer (*Cervus elaphus*), wild boar (*Sus scrofa*), caribou

(*Rangifer tarandus*), chamois (*Rupicapra* spp.), and ibex (*Capra* spp.), with mountain ungulates grouped at the genus level. These taxa were used to derive taxon-specific habitat suitability and resistance layers, which were later summarized into generalized herbivore and carnivore archetypes. The aim was not to predict movement for any single species in a strict mechanistic sense, but rather to capture broad contrasts in permeability and movement opportunity among ecologically distinct mammalian groups at continental extent.

#### Species distribution modelling

To represent broad-scale habitat suitability and landscape permeability, we developed continental-scale species distribution models (SDMs) for each focal taxon. Using presence-only detection data, we fitted generalized linear mixed models with covariates describing proportional land cover, elevation, and climatic conditions. We then conducted model averaging across all top-supported models with $\Delta$AICc < 8, producing continuous suitability surfaces for each taxon across Europe. These suitability surfaces were interpreted as broad-scale indicators of relative permeability rather than as direct predictions of occupancy or realized dispersal. To convert habitat suitability into resistance, we inverted the final suitability surfaces so that areas of high suitability corresponded to low resistance and areas of low suitability corresponded to high resistance. This step allowed SDM outputs to be integrated with the energetics layer in a common resistance framework.

#### Energetic cost of movement

We quantified the energetic cost of terrestrial movement within a broader resistance framework designed to assess whether climate-driven redistribution among protected areas is both structurally and physiologically feasible (Fig. S3). Energetic travel costs were estimated with the R package enerscape (Berti et al., 2022), which implements Pontzer's first-principles allometric cost-of-transport model (Pontzer, 2016), where

$$E(m,\theta) = 8m^{0.66} + 50(1 + sin(2\theta - 74))m^{0.88}$$

To generalize this formulation across taxa, we used its normalized form:

$$\bar{\bar{E}} = \frac{E(m,\theta) - min_\theta E(m,\theta)}{max_\theta E(m,\theta) - min_\theta E(m,\theta)} = \frac{1 + sin(2\theta - 74)}{2}$$

This yields a unitless index of relative travel cost ranging from 0 to 1 that depends only on slope. Terrain incline was derived at 1-km resolution from the GMTED2010 digital elevation model (Danielson & Gesch, 2011), and travel cost for each focal cell was defined as the maximum energetic cost required to move to one of its four cardinal neighbours, producing a continuous surface of topographic movement cost. This energetics layer was then integrated with habitat-based resistance derived from SDM surfaces, while additional high-resistance features not fully captured by SDMs, including major rivers and lakes, glaciers, motorways, and border barriers, were burned into the resistance layers to improve landscape realism. The slope-derived energetics surface and the SDM-derived resistance surface were then rescaled to a common range of 1–1000 and combined pixel by pixel using the maximum value, such that each cell reflected the dominant constraint on movement, whether topographic energetic cost, habitat-associated resistance, or an added barrier. The resulting composite resistance surface therefore distinguishes landscapes that are structurally connected from those that are also energetically and functionally traversable.

**Figure S2**. Spatial distribution of terrain-based energy costs across Europe, expressed as normalized travel cost (kcal $km^{-1}$) calculated with enerscape from terrain incline at 1-km resolution. Warmer colours indicate higher energetic costs of movement, while cooler colours indicate lower costs.

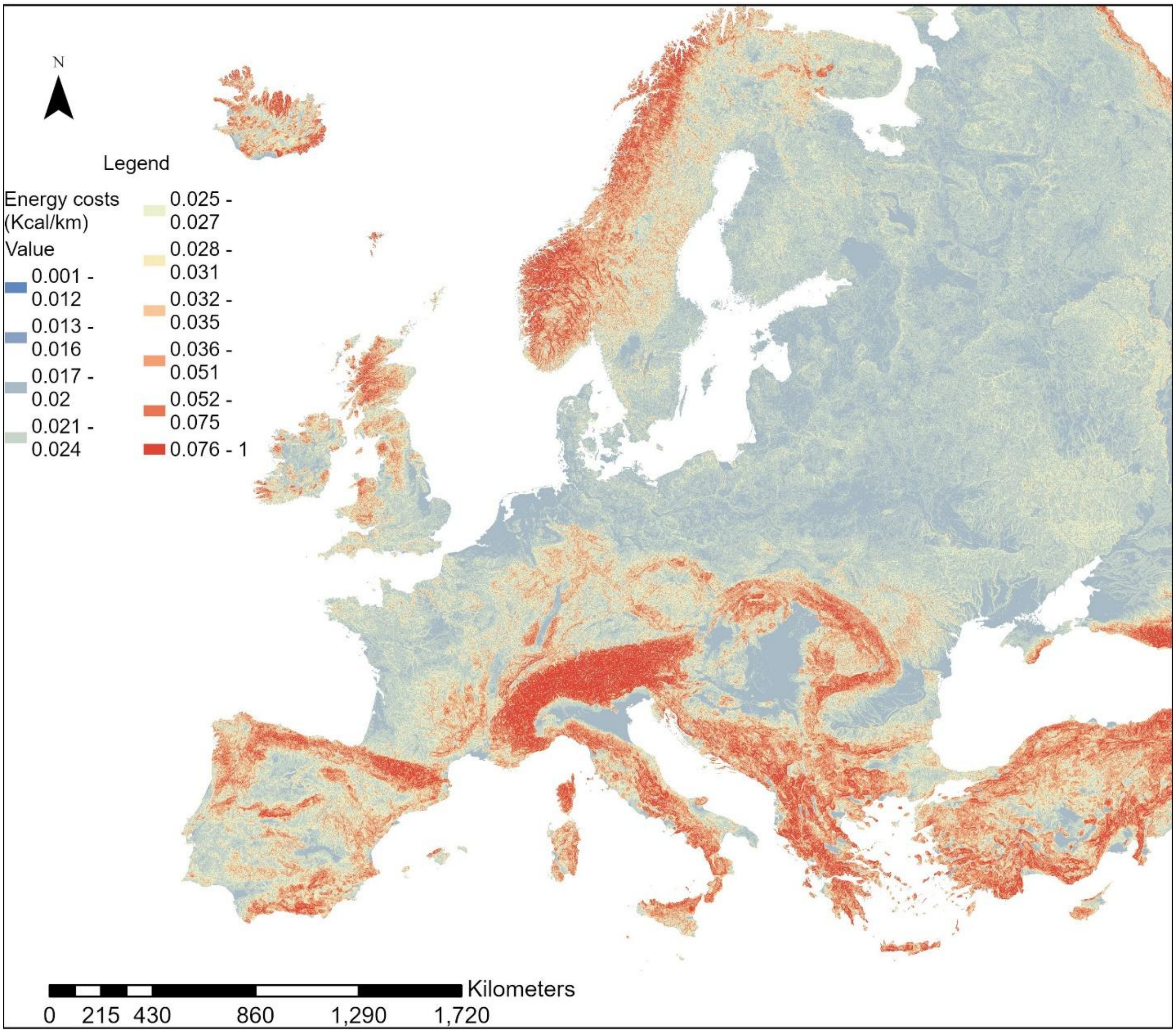

N
Legend
Energy costs (Kcal/km)
Value
0.001 - 0.012
0.013 - 0.016
0.017 - 0.02
0.021 - 0.024
0.025 - 0.027
0.028 - 0.031
0.032 - 0.035
0.036 - 0.051
0.052 - 0.075
0.076 - 1
Kilometers
0 215 430 860 1,290 1,720

### Analytical Overview

Our study combined climate exposure, protected-area geometry, movement energetics, and landscape resistance to assess the feasibility of climate-tracking movement across the European protected-area network (Fig. S3). The workflow comprised four linked components. First, we quantified local climate velocity across terrestrial Europe to estimate the rate at which climatic conditions are expected to shift geographically. Second, we combined climate velocity with protected-area size to derive climatic residence time, a metric describing the expected duration over which a protected area retains its current climate. Third, we constructed energetics-informed resistance surfaces by integrating a slope-based movement-cost layer with habitat-based resistance derived from species distribution models and additional high-resistance barrier features. Finally, we used omnidirectional circuit-theory modelling to estimate cumulative current density and summarize the relative feasibility of movement in and around protected areas.

**Figure S3.** Workflow for deriving energy-based, climate-resilient functional connectivity among European protected areas. Climate change velocity (A) and protected-area geometry (B) are first combined to estimate climate residence time, a protected-area–level metric of climatic persistence (C). This climate residence metric is then integrated with habitat suitability derived from species distribution models (D) and movement energetics (E) to construct energy-based resistance surfaces representing the feasibility of movement across protected areas under climate change. These resistance surfaces are subsequently used in functional connectivity modelling using current-density approaches to estimate potential movement pathways under energetic constraints (F). The resulting connectivity outputs constitute energy-based functional connectivity maps for herbivore and carnivore archetypes across Europe.

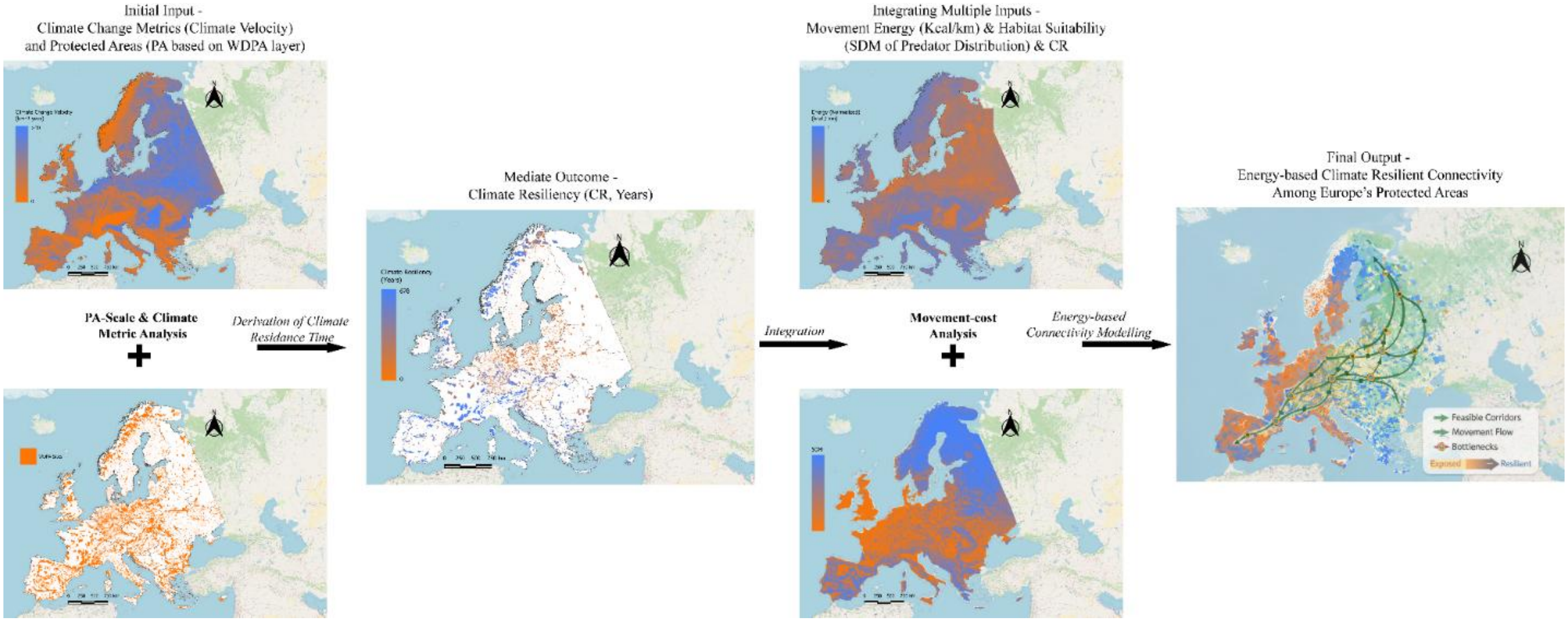
Initial Input -
Climate Change Metrics (Climate Velocity)
and Protected Areas (PA based on WDPA layer)
PA-Scale & Climate
Metric Analysis
Derivation of Climate
Residance Time
Mediate Outcome -
Climate Resiliency (CR, Years)
Integration
Integrating Multiple Inputs -
Movement Energy (Kcal/km) & Habitat Suitability
(SDM of Predator Distribution) & CR
Movement-cost
Analysis
Energy-based
Connectivity Modelling
Final Output -
Energy-based Climate Resilient Connectivity
Among Europe's Protected Areas
Feasible Corridors
Movement Flow
Bottlenecks
Exposed
Resilient
